# Genetic and phenotypic variation between *Tetranychus ludeni* populations from Benelux

**DOI:** 10.1101/2025.05.23.655751

**Authors:** Merijn R. Kant, Ernesto Villacis-Perez

## Abstract

The occurrence of the bean spider mite, *Tetranychus ludeni*, in The Netherlands and Belgium is reported here for the first time. Mite populations were collected from the field and established in the laboratory. Morphological and genetic analyses were performed to determine the species identity of multiple field-collected individuals. Comparing the sequence of the mitochondrial cytochrome oxidase subunit-1 (CO1) of the field populations with sequences available in NCBI revealed the extent of genetic differentiation between *T. ludeni* populations from around the world. The reproductive performance of mated females was assessed on various host plant species relevant to agriculture, including several leguminous species and tomato. These findings highlight the potential of *T. ludeni* to become an agricultural pest in Europe, particularly in the context of increasing global temperatures and agricultural practices.

## Introduction

Phytophagous mites are prevalent across land ecosystems. Mites of the family Tetranychidae (commonly known as spider mites for their ability to spin web) are particularly important for agriculture, with over a hundred species considered pests and at least 10 species classified as major pests^1^. As a group, the Tetranychidae colonise a wide range of host plants from different families, and specialists as well as extremely generalist species have been reported^2^. Specialist species, such as the tomato specialist *Tetranychus evansi*^3^ or the grass specialist *Oligonychus pretensis*^4^ have evolved traits that allow them to efficiently exploit their host plants. In contrast, the ability of generalist species, such as *Tetranychus urticae*, to colonise many different plant species is attributed to a large repertoire of mechanisms that allow it to deal with a diverse array of plant defences^5^. The diversity of adaptive mechanisms found across and within species is likely a result of the on-going arms race between mites and their hosts. For example, patterns of genetic variation associated with host plant adaptation are persistent across space and time in populations of *T. urticae* found in the Dutch dune ecosystem^*6*^.

Among the physiological traits that make spider mites particularly successful pests is their short life cycles and high reproductive capacity^2^. Increasing temperatures and relatively dry conditions can lead to larger reproductive output and faster development, making spider mite species that do not have an important economic impact at the moment, a potential risk in a warmer future^7,8^. *Tetranychus ludeni*, known as the bean spider mite, has a wide geographical distribution, although it is most common in tropical and sub-tropical regions^9^. *T. ludeni* has been reported as a serious pest of bean, soybean, aubergine, sweet potatoes, cucurbitaceous plants, and strawberries^9-14^. *T. ludeni* has the potential of becoming a major pest in agriculture as temperature increases, with a capacity for population growth rates that outmatch that of *T. urticae*^*7*^. Furthermore, using tomato as an experimental host, it has been suggested that *T. ludeni* can manipulate plant defences to the benefit of con-specifics, as well as hetero-specific herbivores^15^. This evidence suggests that *T. ludeni* might become a novel major agricultural pest in temperate regions as temperature rises, particularly for leguminous plant species.

Here, the occurrence of two distinct populations of *T. ludeni* from The Netherlands and Belgium is reported, countries where the species had not been found in the field before^1,16^. Morphological and genetic analyses were conducted to assess the species identity of field-collected samples of the two populations. The reproductive performance of these populations was assessed under laboratory conditions on several host plant species as a proxy of their host range.

## Materials and Methods

### Mites and plants

The *T. ludeni* ‘Aub’ population was collected from aubergine (*Solanum melongena*) in the region of Bredene, Belgium (51°14'48.0"N, 2°57'50.2"E) in 2022. A laboratory population was established by isolating a single virgin female during the last juvenile moulting stage from the aubergine leaves, and allowed it to lay eggs on a detached common bean (*Phaseolus vulgaris*) leaf. Spider mites are haplodiploid, and thus unfertilised eggs develop into haploid males and fertilised eggs develop into diploid females^2^. After male offspring developed into adults (approximately 10-12 days in this species), the mother was isolated with one of her sons to mate on a new leaf, and this process was conducted for five generations of mother-son matings. The population was expanded and maintained on detached bean leaves and kept at standard conditions (25C, 60% relative humidity, 16:8hours light:dark photoperiod).

The *T. ludeni* ‘Hans’ population was collected from common bean in The Netherlands (52°41'51"N, 5°14'23"E) in 2024. The bean leaves were infested simultaneously with multiple mite species including *T. ludeni, T. urticae* green and red forms, as well as several predatory mites. Fifteen *T. ludeni* females were picked manually from the population with a soft brush and each one was isolated to a common bean leaf disc (15mm diameter). After four days, each female was collected separately for DNA extraction. Each mite was crushed in a mixture of 20ul STE buffer + 2ul proteinase K using a pipette tip, followed by incubation for 30 minutes at 37C, 7 minutes at 95C and immediately stored at -20C until further analyses. The same procedure was conducted with 15 Aub females.

Seeds of four leguminous plant species, i.e. common bean (*Phaseolus vulgaris* cv. DWZR), lima bean (*Phaseolus lunatus* cv. Pole lima bean), broad bean (*Vicia faba* cv. Witkiem), soybean (*Glycine max* cv. Chiba green), and of three tomato (*Solanum lycopersicum*) cultivars (cv. Moneymaker, cv. Castlemart and cv. Castlemart *def-1*) were sown in Jongkind soil N.3 (Jongkind B.V., The Netherlands) and grown at 25C, 60-70% RH and with a L16:D8 photoperiod in the greenhouse at the University of Amsterdam. Leguminous species were two weeks old and tomatoes were three weeks old at the time of the experiments.

### Species identification

Morphological and genetic analyses were conducted in parallel to determine the species identity of *T. ludeni* individuals. Morphological analyses were conducted blindly, i.e., before their genetic identities had been determined. Twenty adult female and 20 adult male individuals from the Aub and Hans populations were collected separately in 70% ethanol before subsequent analyses. Individuals were first cleared in lactic acid for 24 hours and then mounted in modified Hoyer’s medium on microscope slides, as described previously^17^. Males were positioned laterally and females with dorsum up in order to identify discriminating characters, such as the angulation of the knob in the male aedeagus and the alignment of proximal tactile setae in tarsus I with the proximal pair of duplex setae in females.

Genetic identification was conducted by amplifying and sequencing the so-called Folmer fragment within the mitochondrially-encoded cytochrome oxidase subunit 1 gene (CO1), using primers LCO1490 (5'-ggtcaacaaatcataaagatattgg-3') and HC02198 (5'-taaacttcagggtgaccaaaaaatca-3')^18^. DNA from the 15 individual females from the Hans and Aub populations was used as template for CO1 amplification in a 25uL PCR reaction, with 0.2uL DreamTaq polymerase (5U), 0.4uL forward and reverse primers [10uM], 5uL dNTPs [1mM], 3.75uL of 10X DreamTaq buffer, 1.25uL of BSA, 11uL H_2_O and 3uL of DNA template. The reaction included an initial denaturation step at 94C for 4min, followed by 35 cycles of 94C x 1min, 48C x 1min, 72C x 1min, and a final step at 72C x 4min. PCR products were checked visually on a 1% agarose gel to confirm the presence of the expected product of ∼700bp. Sanger sequencing of the PCR products was conducted by Macrogen, The Netherlands, using separate reactions for forward and reverse primers. The quality of the sequences was assessed by inspecting the chromatograms visually using SnapGene software (www.snapgene.com) and sequences trimmed at the edges were submitted as queries against the NCBI database.

### *T. ludeni* performance across host species

The reproductive performance of adult females was quantified to estimate the fitness of the *T. ludeni* populations across the four leguminous species and the three tomato cultivars. Performance experiments were conducted using age-synchronised females, which were obtained by transferring approximately 150 mated females from each mite population to a detached common bean leaf for two days and then removing the adults to let the eggs develop to adulthood as a cohort for 12 days. Individuals used for experiments were 2±1 days old.

Age-synchronised females from the Hans and Aub populations were transferred individually to a leaf disc (15mm diameter) of each host plant species, surrounded by wet cotton wool. Female survival was assessed daily for a total of four days. Individuals were scored with a 1 if alive, with a 0 if dead, and with a 0.5 when the mite got stuck to the cotton wool, in which case it was brought back to the centre of the disc. The total number of eggs laid per female was scored at day four, and the data was corrected for female mortality. Only females that survived for at least two days were included in the analyses, thus n = 24 – 32 individuals per population. Differences in reproductive performance where first analysed in an omnibus linear model (*aov* in R v4.4.2) with population, plant species and their interaction as fixed effects. Separate linear models were fit to the subsets of data per host plant species with population set as fixed effect, or with mite population as fixed effect. Model fit was evaluated by assessing the normality and heteroscedasticity of the residuals. Data were square root-transformed to fit model assumptions when needed.

## Results and Discussion

### Morphological and genetic identification of *Tetranychus ludeni*

Male and female individual mites were examined for morphological characters commonly used to determine species identity between spider mite species, without any prior knowledge of their genetic identity. Discriminating characters are only distinguishable with an appropriate mounting procedure, particularly in the shape of the male aedeagus, and thus securing enough individuals that are properly mounted is imperative to identify species in the genus *Tetranychus*^17^. We analysed at least 10 individuals per sex that were correctly mounted, and observed that these characters varied between individuals to some extent, which is not an uncommon phenomenon in spider mites (e.g.^19^). Nonetheless, based on the absence of the posterior angulation of the knob of males and on the alignment of setae in females, all individuals of the Hans and the Aub populations were deemed to belong to *Tetranychus ludeni*.

To assess the species identity and the extent of genetic differentiation between the Hans and Aub populations, a ∼700bp stretch of the CO1 gene from multiple individual females was sequenced. One single CO1 haplotype was found across all individuals within each population. This was expected for the inbred Aub population. However, all individuals from the Hans population, which were collected directly from the field in The Netherlands, also harboured only a single CO1 haplotype. The sequences from the Aub and Hans populations will be deposited into NCBI upon peer-reviewed publication. The CO1 sequences were used as queries in BLASTn searches against the NCBI databank. The best hit for both queries was an accession with the entire mitochondrion genome of *Tetranychus ludeni* (GenBank: KJ729018.1) reported from China^20^, followed by sequences belonging to *T. ludeni* populations from Australia (e.g., GenBank: KX281712.1). These results support the identification of Hans and Aub individuals as *T. ludeni* inferred from the morphological analyses, thus constituting robust evidence for the occurrence of *T. ludeni* in The Netherlands and in Belgium, places where the species has not been reported previously^1,16^, despite being considered a species native to Europe^1^.

A search in GenBank with the query ‘Tetranychus ludeni’ (performed on 5^th^ May 2025) resulted in multiple hits from nuclear and mitochondrial genes. The CO1 sequences available from Germany (MZ151410.1), Australia (KX281712.1), Japan (AB736051.1), India (JX075252) and the USA (MW326486) were aligned to the complete *T. ludeni* mitochondrial genome from China (KJ729018.1), along with the CO1 sequences from Aub and Hans. Genetic differentiation was assessed along a 631bp stretch for most samples, except for the sequences from Japan, the USA and India, which only overlapped in 300-400bp with the sequences obtained from this study due to different primers used to target the CO1 gene (Figure 1). Interestingly, the CO1 of the Aub population resembled all other available *T. ludeni* CO1 haplotypes closely, with only one indel found at the edge of the sequence when compared to the reference mitochondrial genome. In contrast, the Hans population differed greatly from the other CO1 sequences, with variants identified in 54 positions along 631bp when compared to the reference *T. ludeni* mitochondrial genome. The sequence from the USA also differed from the reference, but only in 5 positions in a 408bp stretch. Thus, based on CO1 sequences, it seems that there is distinct genetic variation across *T. ludeni* populations, with at least three different haplotypes found across different continents, including Europe, Australia, North America and Asia.

**Figure 1.**
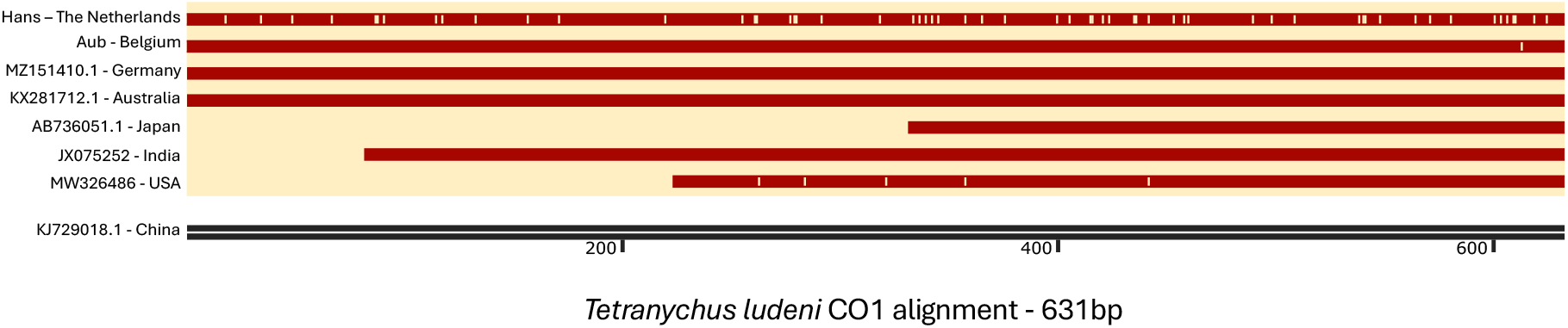
Alignment of CO1 sequences from *Tetranychus ludeni* populations. Sequences obtained from individuals from the Hans and Aub populations, collected in The Netherlands and Belgium respectively, were aligned together with the available CO1 sequences deposited in NCBI (red lines) to the complete mitochondrial genome of *T. ludeni* from a population from China (black double line, bottom)^20^. Positions that differ from the reference sequence are highlighted along the red lines with yellow ticks.

### Variation in host range between *T. ludeni* populations

The genetic differences found between the Aub and Hans populations motivated an analysis of their phenotypic differences. We quantified the reproductive performance of adult females as a proxy of their host range. The reproductive performance of adult females was assessed on four leguminous hosts and on three tomato varieties, among which cv. Castlemart *def-1*. This variety is a tomato mutant with impaired jasmonic acid-related defences^21^, and comparing it to the wildtype (cv. Castlemart) background thus reveals whether mites are resistant or susceptible to this class of tomato defences. There was a statistically significant interaction between *T. ludeni* population and plant species (F_6, 399_ = 20.24, p < 0.001), indicating that reproductive performance of each population depends on the plant species tested. Overall, the reproductive performance of both populations was 3 to 4-fold higher on leguminous hosts than on tomato (Figure 2).

**Figure 2.**
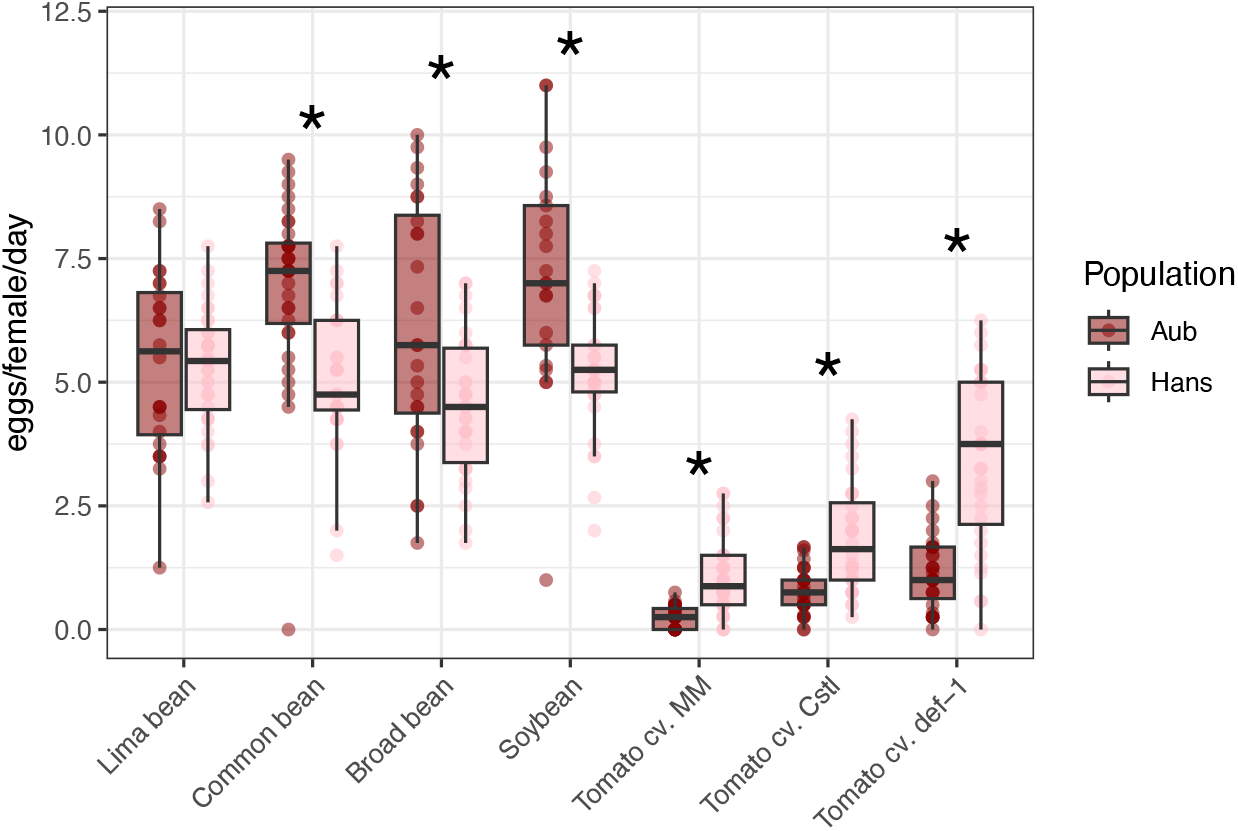
Reproductive performance of *Tetranychus ludeni* across different host plant species. The number of eggs laid per female per day (y-axis) of age-synchronised individuals from the two *T. ludeni* populations, Aub and Hans (legend), on different host species (x-axis). Boxplots show the individual data points, the median (black line within the interquartile box), and the bottom and top 25% quartiles of the data values (whiskers). Asterisks above a pair of boxplots indicate a significant difference according to independent linear models fitted within a plant species, between populations. MM = Moneymaker; Cstl = Castlemart.

The differences in performance between the two populations were compared within host using independent linear models. Interestingly, the performance of Aub was higher than the performance of Hans on all leguminous hosts except for lima bean, and it produced the highest number of eggs laid on soybean. In contrast, Hans had a significantly higher performance on the three tomato hosts than Aub (Figure 2). Since Aub was originally sampled from aubergine this finding was unexpected, as this species belongs to the Solanaceae family, like tomato. The low performance of Aub on tomato could be potentially explained by the fact that it is an experimentally inbred population, and thus could have suffered from inbreeding depression expressing as a decrease in fitness. However, its performance on leguminous hosts was high, suggesting that this was not the case. It is possible that, since both populations were reared on bean in the laboratory, they accepted leguminous hosts more readily than tomato. Another non-mutually exclusive possibility is that inbreeding Aub on bean resulted in the selection of alleles determining their acceptance of leguminous plant species as food at the expense of alleles that determine acceptance of tomato.

Comparing the performance on Castlemart vs. *def-1* of Aub (F_1, 61_ = 2.95, p = 0.09) and of Hans individuals (F_1, 62_ = 16.28, p < 0.001) revealed differences in how they cope with tomato as a host. The performance of Aub was very low and not significantly different between the *def-1* tomato and the corresponding Castlemart wildtype, although a feeble trend towards a higher oviposition on *def-1* was observed (Figure 2). This indicates Aub struggles with tomato in general and not only the traits related to jasmonate defences. In contrast, the performance of Hans was also low on tomato but increased significantly on the jasmonate mutant compared to the wildtype, indicating that the performance of Hans, unlike that of Aub, is negatively affected by jasmonate defences. These findings suggest that there is distinct intraspecific variation in the mechanisms of host use within *T. ludeni*. (Figure 2).

### Implications of variation in host use by *T. ludeni* for agriculture

The set of host plant species chosen to assess mite-host compatibility in this study represent two key agricultural species. Tomato is an economically important European crop, with ∼6.7 million tons produced in the continent in 2024, out of which ∼1-1.5 million tons were produced separately in Spain, The Netherlands, Italy and Poland, in 2024^22^. *T. ludeni* has been found on commercial tomato^12^ and is able to suppress tomato defences, thereby making the plant susceptible to subsequent pests and diseases^15^, similar to its sister species, the tomato pest *T. evansi*^1,23^, as well as some ecotypes of *T. urticae*^*24,25*^. *T. evansi* produces an immense amount of silk web around its host plant, hampering the effectiveness of predators to access individuals in the population, as well as access of competitors to resources^2,26^. Visual inspection of plants infested with *T. ludeni* suggest that this species also produces large amounts of dense web, but whether this differs between populations, or whether this hampers competitors and predators to a similar extent as *T. evansi* remains to be determined. In addition, whether *T. ludeni* and *T. evansi* can hybridise is not known, but given their close genetic relatedness^27^, it is worth investigating whether alleles related to host utilization can be exchanged between these two species.

In Europe, legumes are not produced to the same scale as tomatoes and their production comes largely from the American continent. For example, around 95% of the soybean consumed in the EU is imported, mainly from the United States (∼5.2 million tons in 2018) and Brazil (∼1.3 million tons in 2018)^28^. The European Commission has identified a major gap between the amount of protein produced and the amount imported to the EU, with over 70% of protein-rich crops being imported in 2018^29^. Recently, the EU launched a series of plans to boost legume production in the continent^30^. Soybean cultivation in Europe likely requires modified varieties in order to enable large-scale cultivation under the European biotic and abiotic environmental conditions^31^. Among the traits needed for successful soybean cultivation in Europe, increasing the resistance to crop pests and pathogens while securing a high yield is a major challenge^32^. *T. ludeni* is a common species associated with soybean cultivation in Brazil^11^, and even though moderate mite control was observed on genetically modified soybean cultivars^33^, the data presented here suggest that intraspecific variation within *T. ludeni* populations may lead to rapid adaptations, as in *T. urticae*^*34*^. Furthermore, increasing temperatures impact life-history traits and reproductive strategies in *T. ludeni*, facilitating range expansions and its potential of becoming a major pest in agriculture, to levels comparable to the cosmopolitan pest *T. urticae*^7,35^.

### Future perspectives

The genetic and phenotypic differences related to host use observed between the two *T. ludeni* populations in this study bring forth the question of whether there is a genetic, heritable basis that determines its performance on host plants. If that is the case, the differences in performance on legumes vs. tomatoes could be used to map genetic loci associated with these differences, as it has been done previously for *T. urticae*^*36*^. Future work will focus on assessing whether *T. ludeni* populations are reproductively compatible with each other, and if that is the case, hybrids can be used to elucidate the genetic basis of host plant use. In addition, this study confirms a recurrent observation: *T. ludeni* is often found in mixed populations, where other herbivorous mites reside on the same leaf as *T. ludeni* (e.g.^9,37^). This raises the question of how different competitor mite species interact, either directly or indirectly via host plants, and whether *T. ludeni* facilitates, or restricts, host colonisation for competitors.

## Acknowledgements

We would like to thank Dr. N. Wybouw and Dr. F. Faraji for their assistance with field sampling and morphological analyses, and for reading early versions of the manuscript. This work was supported by the Dutch Research Council (NWO-VICI 19391 to MRK).

## References

1 Migeon, A., Nouguier, E. & Dorkeld, F. in Trends in Acarology 557–560 (Springer, 2024).

2 Helle, W. & Sabelis, M. W. Spider mites: their biology, natural enemies and control. (Elsevier Science Publishers B.V., 1985).

3 Bruinsma, K. et al. Host adaptation and specialization in Tetranychidae mites. Plant Physiology 193, 2605–2621 (2023). 10.1093/plphys/kiad412

4 Bui, H. et al. Generalist and specialist mite herbivores induce similar defense responses in maize and barley but differ in susceptibility to benzoxazinoids. Frontiers in Plant Science 9, 1222 (2018).

5 Grbic, M. et al. The genome of Tetranychus urticae reveals herbivorous pest adaptations. Nature 479, 487–492 (2011). https://doi.org:http://www.nature.com/nature/journal/v479/n7374/abs/nature10640.html#supplementary-information

6 Villacis-Perez, E. et al. Adaptive divergence and post-zygotic barriers to gene flow between sympatric populations of a herbivorous mite. Communications Biology 4, 853 (2021). 10.1038/s42003-021-02380-y

7 Gotoh, T., Moriya, D. & Nachman, G. Development and reproduction of five Tetranychus species (Acari: Tetranychidae): Do they all have the potential to become major pests? Experimental and Applied Acarology 66, 453–479 (2015). 10.1007/s10493-015-9919-y

8 Teodoro-Paulo, J. et al. Rising temperatures favour defence-suppressing herbivores. Journal of Pest Science, 1–14 (2024).

9 Zhang, Z.-Q. Taxonomy of Tetranychus ludeni (Acari: Tetranychidae) in New Zealand and its ecology on Sechium edule. New Zealand Entomologist 25, 27–34 (2002). 10.1080/00779962.2002.9722091

10 Maric, I., Međo, I. & Marčić, D. The dark-red spider mite, Tetranychus ludeni Zacher (Acari: Tetranychidae)–a new pest in Serbian acarofauna. Pesticides and Phytomedicine/Pesticidi i fitomedicina 37, 85–93 (2022).

11 Reichert, M. B., Schneider, J. R., Wurlitzer, W. B. & Ferla, N. J. Impacts of cultivar and management practices on the diversity and population dynamics of mites in soybean crops. Experimental and Applied Acarology 92, 41–59 (2024). 10.1007/s10493-023-00862-8

12 Singh, V. & Chauhan, U. Seasonal population dynamics of spider mite, Tetranychus ludeni Zacher (Tetranychidae) and associated predatory mite, Neoseiulus sp. nr. neoghanii (Phytoseiidae) on tomato (Solanum lycopersicum L. var. Solan gola: Solanaceae) f. Journal of Biological Control, 37–40 (2018).

13 Castro, B. et al. Preference of red mite Tetranychus ludeni Zacher (Acari: Tetranychidae) to sweet potato genotypes. Brazilian Journal of Biology 79, 208–212 (2018).

14 Reddy, G. V. P. Comparative effectiveness of an integrated pest management system and other control tactics for managing the spider mite Tetranychus ludeni (Acari: Tetranychidae) on eggplant. Experimental & Applied Acarology 25, 985–992 (2001). 10.1023/A:1020661215827

15 P. Godinho, D., Janssen, A., Dias, T., Cruz, C. & Magalhães, S. Down-regulation of plant defence in a resident spider mite species and its effect upon con- and heterospecifics. Oecologia 180, 161–167 (2016). 10.1007/s00442-015-3434-z

16 PESI. Pan-European Species directories Infrastructure, <www.eu-nomen.eu/portal> (2025).

17 Faraji, F. & Bakker, F. A modified method for clearing, staining and mounting plant-inhabiting mites. EJE 105, 793–795 (2013).

18 Folmer, O., Black, M., Hoeh, W., Lutz, R. & Vrijenhoek, R. DNA primers for amplification of mitochondrial cytochrome c oxidase subunit I from diverse metazoan invertebrates. Molecular Marine Biology and Biotechnology 3, 294–299 (1994).

19 Xue, W. X. et al. Incomplete reproductive barriers and genomic differentiation impact the spread of resistance mutations between green-and red-colour morphs of a cosmopolitan mite pest. Molecular Ecology (2023).

20 Chen, D.-S. et al. The Complete Mitochondrial Genomes of Six Species of Tetranychus Provide Insights into the Phylogeny and Evolution of Spider Mites. PLOS ONE 9, e110625 (2014). 10.1371/journal.pone.0110625

21 Howe, G. A., Lightner, J., Browse, J. & Ryan, C. A. An octadecanoid pathway mutant (JL5) of tomato is compromised in signaling for defense against insect attack. The Plant Cell 8, 2067–2077 (1996). 10.1105/tpc.8.11.2067

22 EuropeanCommission. Fresh tomatoes dashboard, <https://agridata.ec.europa.eu/extensions/DashboardTomato/Dashboard.html> (2025).

23 Sarmento, R. A. et al. A herbivore that manipulates plant defence. Ecology Letters 14, 229–236 (2011). 10.1111/j.1461-0248.2010.01575.x

24 Kant, M. R., Sabelis, M. W., Haring, M. A. & Schuurink, R. C. Intraspecific variation in a generalist herbivore accounts for differential induction and impact of host plant defences. Proceedings of the Royal Society B-Biological Sciences 275, 443–452 (2008). 10.1098/rspb.2007.1277

25 Alba2009_Article_ResistanceToTheTwo-spottedSpid.

26 Blaazer, C. J. H. et al. Why do herbivorous mites suppress plant defenses? Frontiers in Plant Science 9, 1057–1057 (2018). 10.3389/fpls.2018.01057

27 Matsuda, T., Kozaki, T., Ishii, K. & Gotoh, T. Phylogeny of the spider mite sub-family Tetranychinae (Acari: Tetranychidae) inferred from RNA-Seq data. PloS one 13, e0203136–e0203136 (2018). 10.1371/journal.pone.0203136

28 Rosario, D. & Robin, C. (ed European Commission) (2019).

29 CropLifeEurope. The EU Protein gap - trade policies and GMOs, <http://www.europabio.org> (2018).

30 EuropeanCommission. Europe Soya Declaration, <https://www.donausoja.org/wp-content/uploads/2022/01/joined-declaration.pdf> (2017).

31 Monfort, M., Buitink, J., Roeber, F. & Nogué, F. Genome editing, an opportunity to revive soybean cultivation in Europe. The Plant Journal 121, e17266 (2025). 10.1111/tpj.17266

32 Gao, M., Hao, Z., Ning, Y. & He, Z. Revisiting growth–defence trade-offs and breeding strategies in crops. Plant Biotechnology Journal 22, 1198–1205 (2024). 10.1111/pbi.14258

33 Rode, P. d. A., Toldi, M., Reichert, M. B., Johann, L. & Ferla, N. J. Development of Tetranychus ludeni (Acari: Tetranychidae) on transgenic soybean cultivars. Phytoparasitica 46, 137–141 (2018). 10.1007/s12600-017-0634-6

34 Shu, Y., Romeis, J. & Meissle, M. No Interactions of Stacked Bt Maize with the Non-target Aphid Rhopalosiphum padi and the Spider Mite Tetranychus urticae. Front Plant Sci 9, 39 (2018). 10.3389/fpls.2018.00039

35 Zhou, P., He, X. Z., Chen, C. & Wang, Q. Reproductive Strategies That May Facilitate Invasion Success: Evidence From a Spider Mite. Journal of Economic Entomology 114, 632–637 (2021). 10.1093/jee/toaa313

36 Villacis-Perez, E. et al. Independent Genetic Mapping Experiments Identify Diverse Molecular Determinants of Host Adaptation in a Generalist Herbivore. Molecular Ecology n/a, e17618 (2024). 10.1111/mec.17618

37 Ragusa, E., Sinacori, M. & Tsolakis, H. First record of Tetranychus ludeni Zacher (Acariformes: Tetranychidae) in Italy. International Journal of Acarology 45, 26–28 (2019).

